# Protective Role of CBD Against Nicotine Pouch–Induced Seizure Aggravation and Alterations in Brain Glymphatic Biomarkers

**DOI:** 10.1101/2025.05.28.656668

**Authors:** Bidhan Bhandari, Sahar Emami Naeini, Hannah M Rogers, Abdullah Hassan Alhashim, Jack C Yu, Mohammad Seyyedi, Nancy Young, Évila Lopes Salles, Lei P Wang, Babak Baban

**Affiliations:** Department of Oral Biology and Diagnostic Sciences, Dental College of Georgia, Augusta University, Augusta, GA, USA; Center for Excellence in Research, Scholarship and Innovation (CERSI), Dental College of Georgia, Augusta, Augusta University, Augusta, GA, USA; The Graduate School, Augusta University, Augusta, GA, USA; Georgia Institute of Cannabis Research, Medicinal Cannabis of Georgia LLC, Augusta, GA, USA; Department of Surgery, Medical College of Georgia, Augusta University, Augusta, GA, USA; Piedmont Ear, Nose, Throat and Related Allergy, Atlanta, GA, USA Medical College of Georgia, Augusta University, Augusta, GA, USA; Department of General Dentistry, Dental College of Georgia, Augusta University, Augusta, GA, USA; Department of Neurology, Medical College of Georgia, Augusta University, Augusta, GA, USA

**Author notes:** **Corresponding authors:** Babak Baban, Ph.D., MPH, MBA, FAHA., Lei P Wang, Ph.D.

## Abstract

Nicotine pouches are increasingly popular as a smokeless alternative to tobacco, yet their long-term neurological effects remain poorly understood. In this preclinical study, we investigated the time-dependent impact of oral nicotine pouch exposure on seizure susceptibility, glymphatic function, and neuroinflammation in mice, and evaluated the therapeutic potential of inhaled cannabidiol (CBD). Using the Racine scale, we found that acute nicotine exposure transiently reduced seizure severity, while chronic exposure significantly exacerbated seizures and impaired glymphatic integrity, as evidenced by downregulation of aquaporin-4 (AQP4). Chronic nicotine also elevated circulating levels of HMGB1 and IL-6, indicating sustained systemic inflammation. Notably, inhaled CBD reversed these pathological changes, reducing seizure severity, restoring AQP4 expression, and normalizing inflammatory markers. Molecular analyses further revealed upregulation of BDNF and c-FOS with chronic nicotine exposure, which was also mitigated by CBD.

In conclusion, these findings suggest that while nicotine pouches may confer short-term neurophysiological modulation, chronic use poses significant risks for seizure vulnerability and glymphatic dysfunction. Inhaled CBD demonstrates strong neuroprotective potential and may serve as a promising therapeutic approach for individuals exposed to prolonged nicotine use.

**Highlights:**

- Chronic exposure to oral nicotine pouches exacerbates seizure severity in a preclinical model.
- Nicotine-induced seizures are associated with elevated neuroinflammation and impaired glymphatic function.
- Inhaled cannabidiol (CBD) reverses nicotine-induced increases in HMGB1, IL-6, and seizure activity.
- CBD restores Aquaporin-4 expression and reestablishes glymphatic integrity disrupted by nicotine.
- Systems biology analysis reveals an IL-6–centered protein network targeted by CBD, offering mechanistic insight into its neuroprotective effects.

## Introduction

Nicotine pouches have gained popularity as an alternative to smoking and smokeless tobacco products, offering a controlled nicotine delivery system through the oral mucosa, where nicotine is absorbed via mucous membranes and enters the bloodstream (Joshua M Jackson et al., 2023; Chris Deery., 2023; Dulyapong Rungraungrayabkul et al., 2024). Unlike smoking, nicotine pouches do not contain the carcinogens associated with tobacco combustion (Erika Grandolfo et al., 2024). While marketed as a safer alternative to traditional smoking, the full extent of their impact on neurological health, cardiovascular function, and overall well-being remains unclear (Wojciech Hajdusianek et al., 2021; E Ulysses Dorotheo et al., 2024). Despite their design to minimize harmful effects compared to smoking, the potential for nicotine pouches to induce adverse health effects, particularly in the brain and nervous system, has not been thoroughly investigated.

Nicotine itself is a potent stimulant with complex effects on the nervous system (Nihaal Singh et al., 2023). Acute nicotine exposure has been shown to have neuroprotective properties in some contexts, enhancing neuronal activity and promoting certain neurotrophic factors (Arrin C Brooks and Brandon J Henderson., 2021). However, chronic nicotine exposure can exacerbate neurological issues, potentially increasing susceptibility to seizures, impairing cognitive function, and inducing neuroinflammation (Omar Hahad et al., 2021). Nicotine pouches, which provide a steady, controlled release of nicotine, present a unique opportunity to explore both short- and long-term neurological effects, as well as their impact on cardiovascular health (Nadja Mallock-Ohnesorg et al., 2024).

Given these uncertainties, we chose to use a mouse model of epilepsy to investigate the neurological effects of nicotine pouches. Epilepsy was selected as a model to assess how nicotine exposure may alter seizure activity, a reflection of broader neurological function. While this model may not represent all potential neurological consequences, it allows us to examine how acute and chronic nicotine exposure may affect brain function and exacerbate seizure activity.

Cannabidiol (CBD), a non-psychoactive compound derived from cannabis, has gained attention for its neuroprotective and anti-seizure properties (Babak Baban et al., 2021; Bidhan Bhandari et al., 2025). Previous studies have demonstrated that inhaled CBD may be more effective in reducing seizures compared to other delivery methods, such as oral or injection forms, due to its rapid onset and targeted action (Bidhan Bhandari et al., 2025). However, its potential to counteract the detrimental effects of nicotine on the nervous system, specifically in terms of seizure exacerbation and neuroinflammation, has not been fully explored.

In this study, we examine the impact of nicotine pouches on seizure severity in a mouse model of epilepsy, focusing on both acute (60 min) and long-term (1 week) exposure. Additionally, we investigate the potential neuroprotective effects of inhaled CBD in mitigating the adverse consequences of nicotine exposure, assessing seizure activity, neuroinflammation, and other markers of neurological health. This research aims to provide a deeper understanding of the unknown neurological and cardiovascular risks associated with nicotine pouch use and explore the therapeutic potential of inhaled CBD for reducing these risks.

## Materials and Methods

### Animals and Ethical Compliance

All experiments were conducted using adult male C57BL/6 mice (8–10 weeks old) obtained from Jackson Laboratories (USA). Mice were housed under standard laboratory conditions with controlled temperature, humidity, and a 12-hour light/dark cycle, with food and water provided ad libitum. All procedures were approved by the Augusta University Institutional Animal Care and Use Committee (Protocol #2011-0062) and performed in accordance with the NIH Guide for the Care and Use of Laboratory Animals.

### Experimental Groups and Treatment Design

Mice were randomly assigned to five groups (n = 5 per group; replicated in five independent cohorts): **1-** Control: Seizure induction without any pretreatment., **2-** Acute Nicotine: Single-dose oral nicotine (60 minutes prior to seizure)., **3-** Chronic Nicotine: Daily oral nicotine for 7 consecutive days prior to seizure induction. **4-** CBD Only: Single-dose CBD inhalation 30 minutes before seizure induction., **5-** Nicotine + CBD: Chronic nicotine for 7 days followed by CBD inhalation prior to seizure.

### Seizure Induction Protocol

Seizures were induced using intraperitoneal (IP) injection of kainic acid (KA, 20 mg/kg; Fisher Scientific) dissolved in sterile phosphate-buffered saline (PBS, pH 7.4), following established protocols as described previously (Bidhan Bhandari et al., 2025). Mice were monitored continuously post-injection, and seizure activity was scored behaviorally using the Racine scale, which classifies seizure severity from stage 1 (facial automatisms) to stage 5 (generalized tonic-clonic seizures). All scoring was performed by blinded observers to minimize bias.

### Nicotine Administration

Nicotine was delivered using ZYN pouches (Philip Morris International). A nicotine dose of 0.5 mg/kg was applied directly to the oral mucosa using a sterile applicator. Mice received either a single dose (acute exposure) or daily doses for one week (chronic exposure), depending on group assignment.

### CBD Inhalation Treatment

Broad-spectrum CBD was administered at a dose of 10 mg per mouse via aerosolized inhalation using a custom-engineered delivery system (ApelinDx, Thriftmaster Holdings Global Bioscience, TX, USA). Inhalation was performed 30 minutes prior to seizure induction, following a previously established protocol (Bidhan Bhandari et al., 2025).

### Immunofluorescence Analysis

Fixed, paraffin-embedded brain tissues were sectioned and labeled with antibodies targeting BDNF (Thermo Fisher, Cat# 500-P84BT), c-FOS (Thermo Fisher, Cat# BS-12911R-A488), and Aquaporin-4 (AQP4) (Thermo Fisher, Cat# ABN91025UG). Sections were counterstained with DAPI and imaged using a Zeiss fluorescence microscope. Quantification of immunoreactivity was performed using ImageJ software (NIH), with regions of interest manually defined in hippocampal and cortical areas.

### Flow Cytometry for Inflammatory Biomarkers

Peripheral whole blood was collected via submandibular bleeding and processed for intracellular staining of HMGB1 and IL-6 as described previously (Babak Baban et al., 2021; Hesam Khodadadi et al., 2020). Cells were fixed/permeabilized using the eBioScience Fix/Perm kit and labeled with fluorophore-conjugated antibodies, HMGB1 (Biolegend, Cat#651404) and IL-6 (Biolegend, Cat#504508). Flow cytometry was conducted using a NovoCyte Quanteon system. Data were analyzed using FlowJo v11, with gating based on standard forward/side scatter and appropriate isotype controls. Marker expression was reported as the percentage of positive events within the lymphocyte gate.

### Bioinformatic Network Analysis

To explore the molecular pathways influenced by nicotine exposure and modulated by CBD treatment, a protein–protein interaction (PPI) network was generated using the STRING database (version 11.0) (Damian Szklarczyk et al., 2023). Interactions were curated based on experimentally validated and predicted functional associations, with a minimum confidence score cutoff set at 0.4. To further characterize the biological significance of the network components, Gene Ontology (GO) enrichment analysis was performed using Metascape and Enrichr, emphasizing pathways relevant to neuroinflammation, neuronal excitability, glymphatic system function, and immune response regulation.

### Statistical Analysis

Racine scores were analyzed using the Kruskal-Wallis H test followed by pairwise Mann–Whitney U tests. Flow cytometry and immunofluorescence quantifications were compared using one-way ANOVA with Tukey’s post hoc test. Significance was set at p < 0.05 for all analyses. Data are presented as mean ± SEM and analyzed using GraphPad Prism 9.

## Results

### Time-Dependent Effects of Nicotine Pouch Exposure on Seizure Susceptibility and Attenuation by CBD Inhalation

Seizure severity was assessed using the Racine scale in mice exposed to either acute (60-minute) or chronic (7-day) oral nicotine pouch treatment (ZYN), followed by kainic acid (KA)-induced seizures. Color-coded Racine scoring over time is illustrated in Figure 1A, and corresponding group-wise averages are quantified in Figure 1B. Mice receiving a single oral dose of nicotine one hour before KA injection exhibited significantly reduced seizure severity compared to the KA-only group (mean Racine score: 2.1 ± 0.3 vs. 3.8 ± 0.4; p < 0.001). This reduction was reflected in lower Racine scores across the 60-minute post-injection window, suggesting that acute nicotine exposure transiently dampens neuronal excitability and confers short-term seizure protection. In contrast, chronic nicotine administration for seven consecutive days led to a pronounced increase in seizure severity (mean score: 4.6 ± 0.2), significantly higher than both the control and acute exposure groups (p < 0.001). This time-dependent escalation highlights a biphasic effect of nicotine, wherein short-term exposure exerts a suppressive effect on seizure activity, but sustained use leads to heightened excitability and seizure vulnerability.

**Figure 1.**
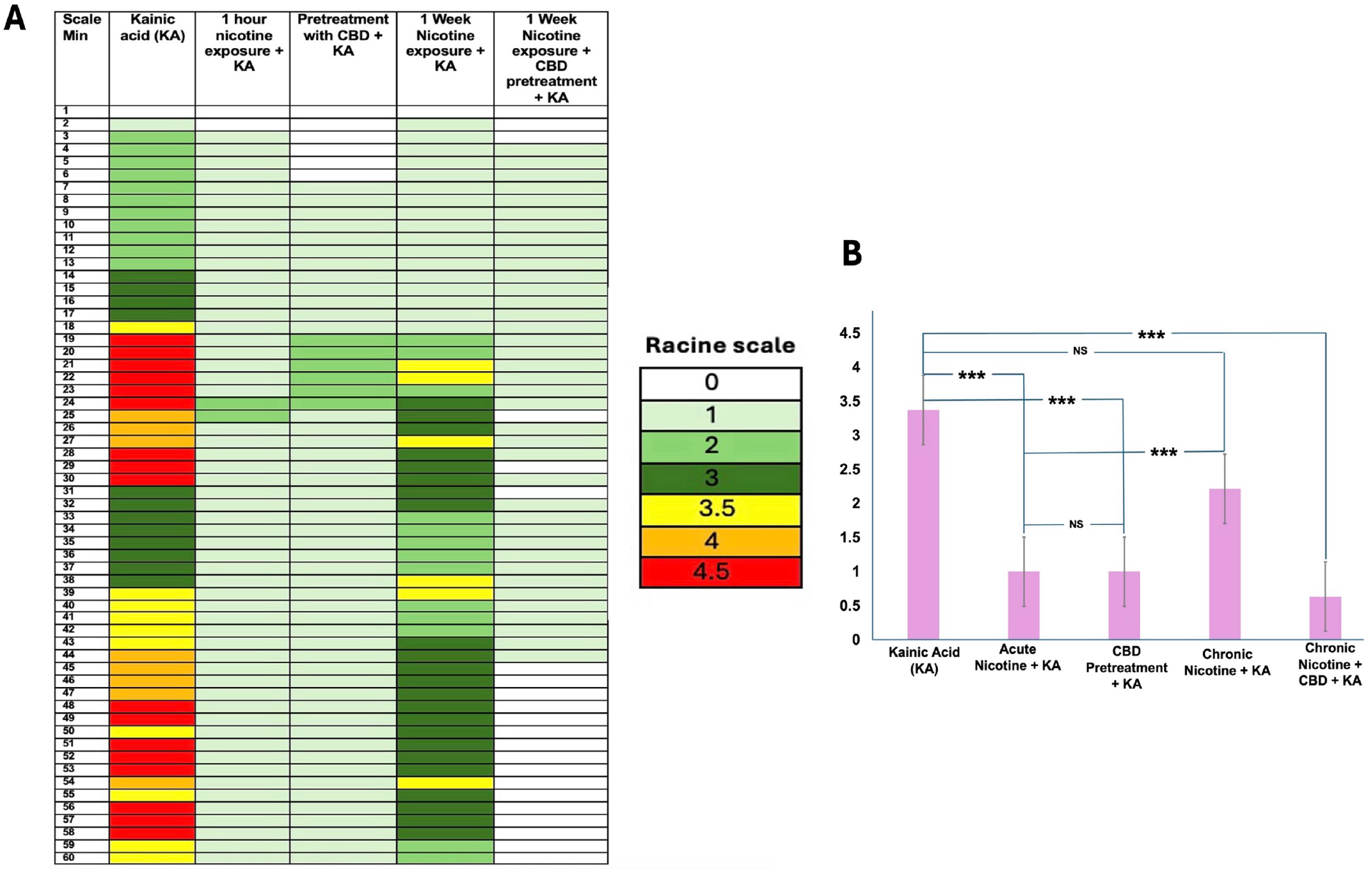
Acute and chronic nicotine exposure differentially modulate seizure severity, with CBD inhalation reversing the proconvulsant effects of chronic nicotine. **(A)** Time-course chart displaying Racine scale scores across 60 minutes for each treatment group. Mice received kainic acid (KA) alone or in combination with nicotine and/or CBD: KA only, acute nicotine exposure + KA, CBD pretreatment + KA, chronic nicotine exposure + KA, and chronic nicotine exposure with CBD pretreatment + KA. Each row represents one minute of post-KA observation. Seizure severity was scored using the Racine scale (0–5), and color-coded for clarity: white (0), light green (1), green (2), dark green (3), yellow (3.5), orange (4), and red (4.5). **(B)** Quantification of average Racine scores for each group. Acute nicotine exposure significantly reduced seizure severity compared to KA alone, while chronic nicotine exposure worsened it. CBD inhalation, both alone and following chronic nicotine, significantly attenuated seizure severity. Statistical analysis was performed using Kruskal–Wallis test followed by Mann–Whitney U test for post hoc comparisons. ***p < 0.001; NS, not significant.

Importantly, mice that received inhaled cannabidiol (CBD) 30 minutes before KA, following one week of chronic nicotine exposure, showed a marked reduction in seizure severity, with Racine scores comparable to those observed in the acute nicotine group (p < 0.001). These findings, also shown in Figure 1B, demonstrate that CBD inhalation effectively reverses the proconvulsant effects of prolonged nicotine exposure. Together, these results support a dynamic interaction between nicotine exposure duration and seizure susceptibility, and identify inhaled CBD as a promising intervention for mitigating nicotine-induced seizure exacerbation.

### Time-Dependent Changes in BDNF and c-FOS Expression are reversed by Inhaled CBD

To investigate the molecular correlates of seizure susceptibility, we assessed hippocampal expression of BDNF, a neurotrophin critical for synaptic plasticity, and c-FOS, an immediate-early gene marker of neuronal activation, using immunofluorescence imaging and quantitative analysis. As shown in Figure 2A, kainic acid (KA) administration markedly increased the co-expression of BDNF and c-FOS compared to sham controls, indicating robust neuronal activation. Acute nicotine exposure (60 min prior to KA) significantly attenuated this effect, with reduced BDNF and c-FOS immunoreactivity, suggesting that short-term nicotine dampens neuronal hyperexcitability (Figure 2B; p < 0.01). In contrast, chronic nicotine exposure over seven days resulted in a pronounced upregulation of both markers, with BDNF and c-FOS expression levels significantly elevated compared to all other groups (p < 0.01), consistent with enhanced neuronal activation and plasticity linked to seizure escalation. Notably, mice pretreated with inhaled CBD (30 minutes prior to KA) following chronic nicotine exposure exhibited a significant reduction in BDNF and c-FOS levels (p < 0.01), restoring expression patterns to levels similar to the acute nicotine and CBD-only groups. These findings demonstrate that chronic nicotine promotes aberrant neuronal activation, while CBD inhalation effectively normalizes this response, reinforcing its neuroprotective potential in the context of sustained nicotine exposure.

**Figure 2.**
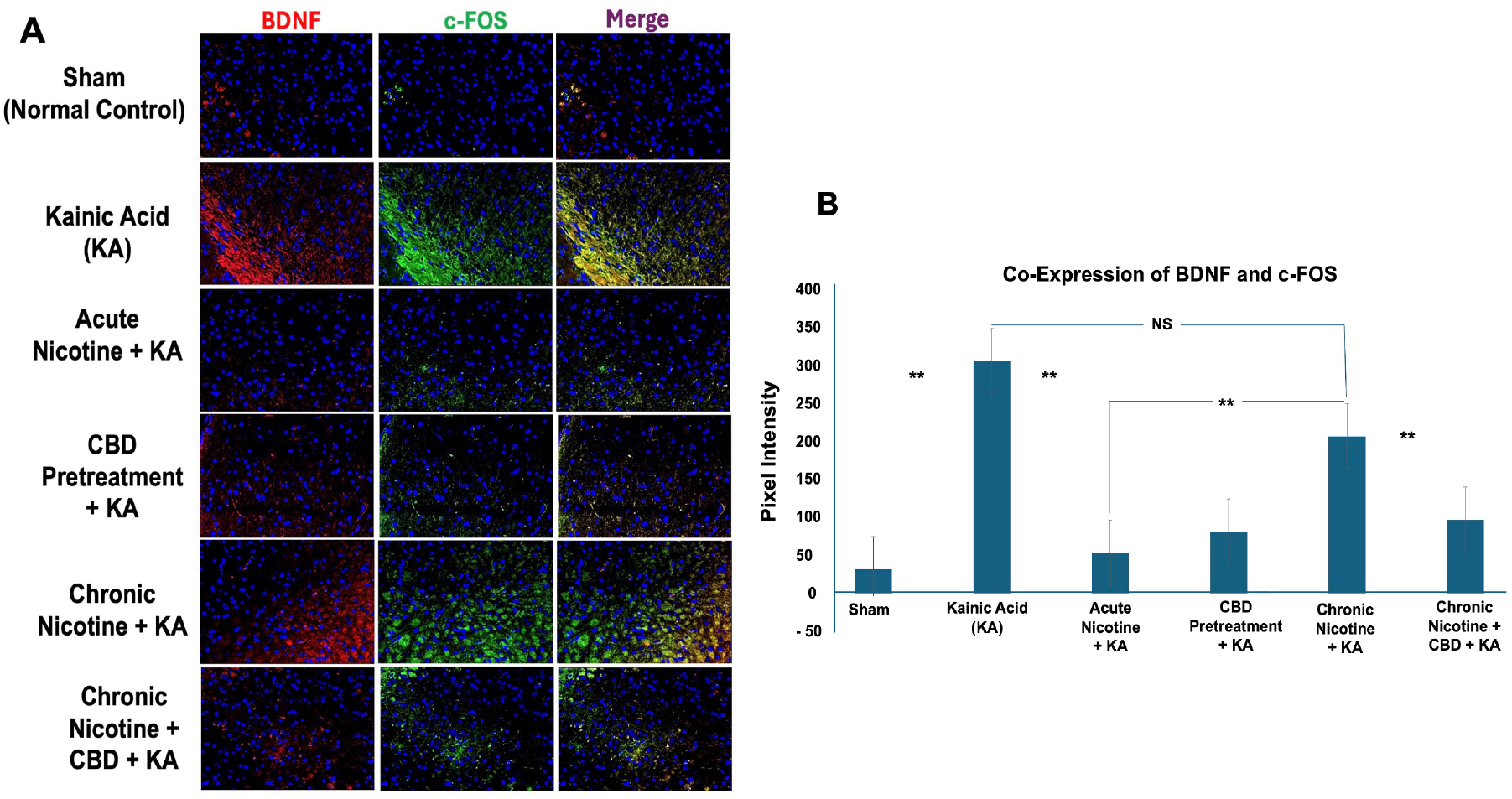
Chronic nicotine exposure amplifies neuronal activity and BDNF signaling, which is reversed by inhaled CBD. **(A)** Representative immunofluorescence images of hippocampal sections showing expression of BDNF (red), c-FOS (green), and merged co-localization (yellow) in the presence of DAPI nuclear stain (blue). Groups include: Sham (untreated control), KA only, acute nicotine + KA, CBD pretreatment + KA, chronic nicotine + KA, and chronic nicotine + CBD + KA. KA exposure led to marked upregulation of BDNF and c-FOS, indicative of heightened excitability and neuronal stress. Acute nicotine and CBD pretreatment attenuated this response, while chronic nicotine exposure further increased co-expression of both markers. Inhaled CBD following chronic nicotine exposure effectively reduced BDNF/c-FOS expression to near-baseline levels. **(B)** Quantification of BDNF and c-FOS co-expression presented as mean pixel intensity. Data show significant elevation in the KA and chronic nicotine groups, and significant reduction with CBD pretreatment or post-chronic exposure (n = 5 per group). Statistical analysis was performed using one-way ANOVA followed by Tukey’s post hoc test. **p < 0.01; NS, not significant.

### Chronic Nicotine Exposure Impairs Glymphatic Function via AQP4 Downregulation, Reversed by Inhaled CBD

To investigate the impact of nicotine and CBD on glymphatic function, we assessed the expression of Aquaporin-4 (AQP4) in the hippocampus using immunofluorescence staining and pixel intensity quantification. As shown in Figure 3A, kainic acid (KA) administration alone led to a marked reduction in AQP4 expression relative to sham controls (p < 0.01), indicating acute disruption of glymphatic homeostasis. Acute nicotine exposure (60 minutes prior to KA) preserved AQP4 expression at levels comparable to controls (NS), suggesting that short-term nicotine use does not compromise glymphatic integrity. In contrast, chronic nicotine exposure for seven consecutive days prior to seizure induction resulted in a substantial downregulation of AQP4 expression (p < 0.01), as visualized by diminished green fluorescence and confirmed in Figure 3B. This finding implicates sustained nicotine use in glymphatic dysfunction, potentially impairing interstitial waste clearance and contributing to neuroinflammatory stress. Notably, mice pretreated with inhaled CBD (30 minutes prior to KA) following chronic nicotine exposure exhibited a significant restoration of AQP4 expression to near-sham levels (p < 0.01), supporting the protective effect of CBD on glymphatic pathways. Together, these results suggest that chronic nicotine disrupts fluid homeostasis within the brain, and that CBD inhalation can effectively restore AQP4 expression and support glymphatic function under neurotoxic conditions.

**Figure 3.**
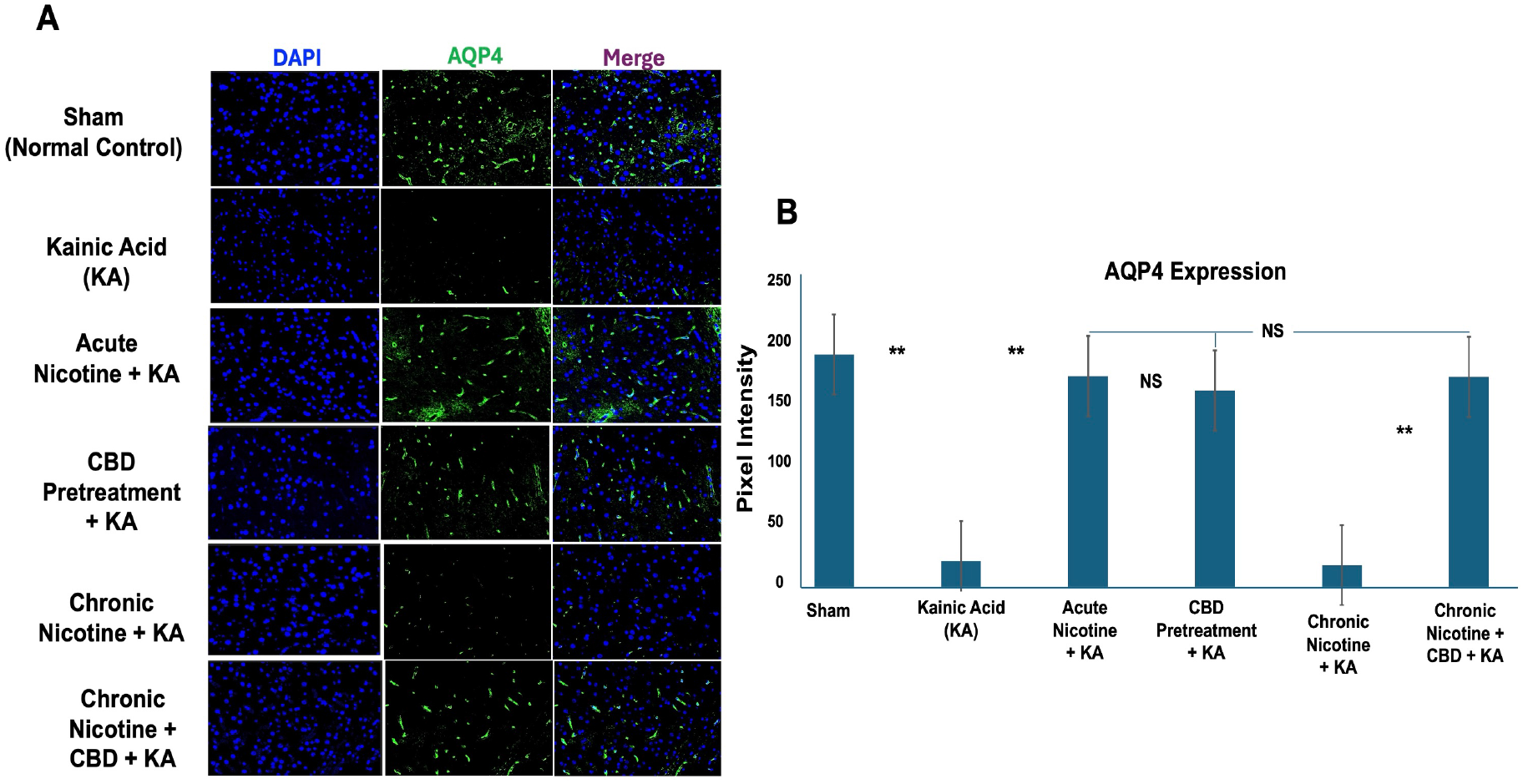
Chronic nicotine exposure suppresses hippocampal AQP4 expression, reversed by inhaled CBD. **(A)** Representative immunofluorescence images showing expression of Aquaporin-4 (AQP4; green) in hippocampal sections, counterstained with DAPI (blue) to label nuclei. Merged panels depict co-localization. Groups include Sham (untreated control), kainic acid (KA), acute nicotine + KA, CBD pretreatment + KA, chronic nicotine + KA, and chronic nicotine + CBD + KA. KA exposure led to a substantial reduction in AQP4 expression, while acute nicotine and CBD pretreatment preserved AQP4 levels. Chronic nicotine exposure further diminished AQP4 signal, consistent with impaired glymphatic function. Notably, CBD inhalation following chronic nicotine exposure restored AQP4 expression to near-control levels. **(B)** Quantitative analysis of AQP4 pixel intensity. KA and chronic nicotine groups showed significant reductions in AQP4 compared to controls (p < 0.01), whereas acute nicotine and CBD-treated groups maintained normal expression. Inhaled CBD reversed chronic nicotine-induced AQP4 loss (p < 0.01). Statistical analysis was performed using one-way ANOVA followed by Tukey’s post hoc test. **p < 0.01; NS, not significant.

### Systemic Inflammation Escalates with Chronic Nicotine Exposure and Is Reversed by Inhaled CBD

To assess the systemic immune response associated with seizure activity and nicotine exposure, we measured the co-expression of IL-6 and HMGB1 in peripheral blood leukocytes using flow cytometry. As shown in Figure 4A, kainic acid (KA) administration led to a significant increase in IL-6 and HMGB1 expression compared to sham controls (p < 0.01), indicating the onset of a systemic inflammatory state. Acute nicotine exposure (60 minutes prior to KA) attenuated this response, resulting in markedly lower IL-6 and HMGB1 levels compared to the KA-only group (p < 0.01), consistent with a short-term anti-inflammatory effect. In contrast, chronic nicotine exposure over seven days prior to KA injection led to a dramatic elevation in the co-expression of IL-6 and HMGB1 (p < 0.01), surpassing the inflammatory response induced by KA alone. This suggests a cumulative, time-dependent exacerbation of systemic inflammation that may contribute to seizure amplification. Remarkably, inhaled CBD, administered 30 minutes prior to seizure induction following chronic nicotine exposure, significantly reduced IL-6 and HMGB1 levels to near-sham values (p < 0.01), as illustrated in Figure 4B. These findings demonstrate that CBD inhalation effectively reverses the proinflammatory milieu induced by prolonged nicotine exposure, supporting its role as a potential therapeutic agent for modulating seizure-associated inflammation.

**Figure 4.**
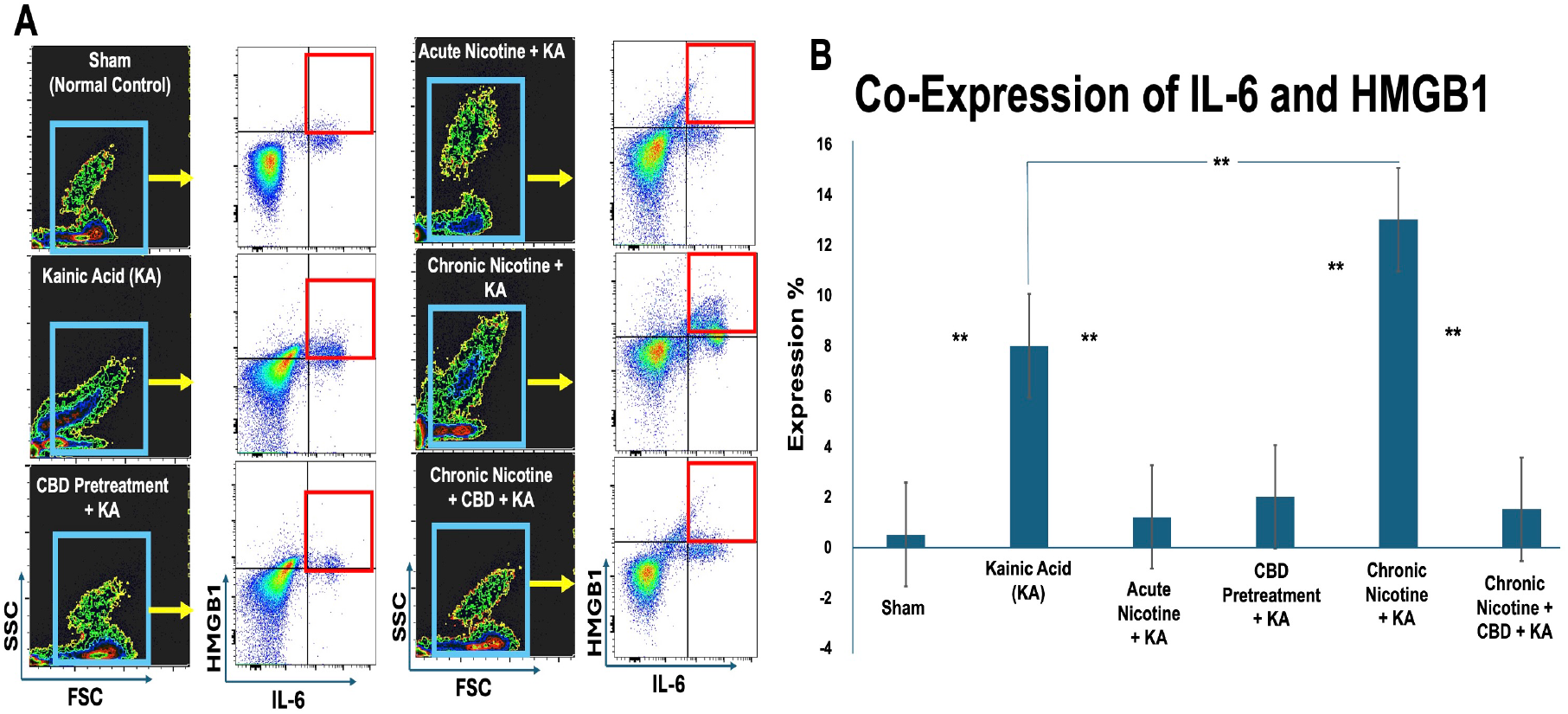
Chronic nicotine exposure amplifies systemic inflammation through IL-6 and HMGB1 co-expression, reversed by inhaled CBD. **(A)** Representative flow cytometry plots showing forward/side scatter (FSC/SSC) gating (left column) and dual-expression plots for interleukin-6 (IL-6) and high-mobility group box 1 (HMGB1) (right column) in peripheral blood leukocytes. Experimental groups include Sham (untreated control), kainic acid (KA) only, acute nicotine + KA, CBD pretreatment + KA, chronic nicotine + KA, and chronic nicotine + CBD + KA. KA and chronic nicotine groups exhibit strong co-expression of IL-6 and HMGB1, while acute nicotine and CBD treatments markedly reduce inflammatory marker expression. **(B)** Quantitative analysis of co-expression percentages for IL-6 and HMGB1. KA significantly elevated co-expression relative to sham controls (p < 0.01), and this effect was further amplified by chronic nicotine exposure (p < 0.01). Acute nicotine and CBD pretreatment alone did not increase inflammatory marker levels, and CBD inhalation following chronic nicotine exposure significantly reduced co-expression back to baseline. Statistical analysis was conducted using one-way ANOVA with Tukey’s post hoc test. **p < 0.01; NS, not significant.

### CBD Rebalances a Nicotine-Responsive Protein Interaction Network Associated with Neuroinflammation and Seizure Vulnerability

To gain mechanistic insight into how nicotine promotes seizure susceptibility and how CBD confers neuroprotection, we conducted a systems-level bioinformatic analysis based on proteins identified in our experimental datasets. Using STRING network modeling, we constructed a protein–protein interaction (PPI) network that revealed a highly interconnected core centered on IL-6, with direct links to key neuroimmune and excitability-associated proteins including HMGB1, FOS, BDNF, S100B, IL1f8, Chrna7, and Lynx1 (Figure 5A). The prominence of IL-6 as a hub node highlights its central regulatory role in mediating the inflammatory and excitatory responses underlying nicotine-induced seizure activity and their reversal by CBD.

**Figure 5.**
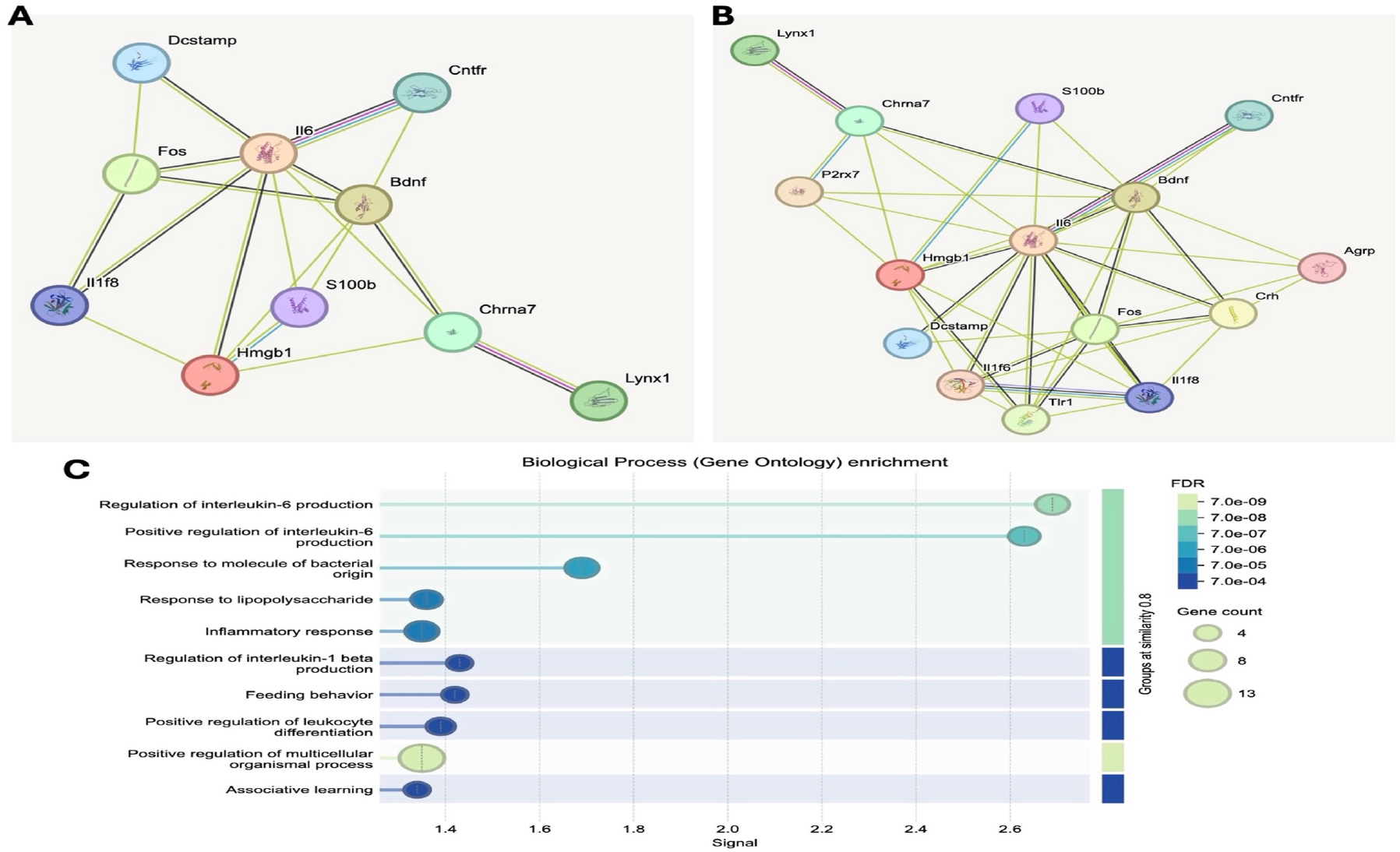
CBD modulates a nicotine-responsive protein network enriched in neuroinflammatory and glymphatic regulatory pathways. **(A–B)** Protein–protein interaction (PPI) networks generated using the STRING v11.0 platform based on experimentally derived and predicted molecular targets from the study. **(A)** A focused network centered around IL-6, showing key connections with neuroinflammatory and neuronal activation markers including BDNF, FOS, HMGB1, IL1f8, and glymphatic-related genes such as S100B and Chrna7. **(B)** An expanded network incorporating additional modulators of neuroimmune signaling, including P2rx7, Crh, Tlr1, Agrp, and others. These networks reflect the integrated roles of cytokine signaling, stress responses, cholinergic tone, and glymphatic activity in nicotine-induced seizure susceptibility and CBD-mediated neuroprotection. **(C)** Gene Ontology (GO) enrichment analysis visualized via bubble plot reveals significantly enriched biological processes, including regulation of IL-6 production, inflammatory response, response to lipopolysaccharide, and leukocyte differentiation, with false discovery rates (FDR) color-coded and gene counts reflected in circle size. Enriched pathways indicate that CBD may exert protective effects by rebalancing immune signaling and excitability within this network.

The expanded PPI network (Figure 5B) incorporated additional stress and immune modulators, including TLR1, Crh, P2rx7, and Agrp, indicating that chronic nicotine exposure disrupts homeostatic control across neuroinflammatory, cholinergic, and hypothalamic-pituitary-adrenal (HPA) stress signaling pathways. These findings are consistent with our flow cytometry and immunostaining data, suggesting that persistent nicotine exposure perturbs molecular circuits that control excitability, immune activation, and glymphatic integrity.

Gene Ontology (GO) enrichment analysis of this network (Figure 5C) revealed significant overrepresentation of biological processes related to regulation of IL-6 production, response to bacterial and lipopolysaccharide stimuli, leukocyte differentiation, and neuroinflammatory response. These enriched pathways closely align with our observed experimental outcomes, particularly the IL-6 and HMGB1 elevations in chronic nicotine-treated animals and their normalization following acute CBD inhalation.

Together, these findings suggest that CBD exerts its neuroprotective effects by restoring balance within a nicotine-disrupted molecular network, with IL-6 acting as a central inflammatory mediator. This systems-level perspective reinforces the therapeutic potential of CBD in mitigating nicotine-related seizure risk and highlights targetable pathways for future intervention strategies.

## Discussion

This study provides compelling preclinical evidence that oral nicotine pouch exposure exerts complex, time-dependent effects on seizure susceptibility, neuroinflammation, and glymphatic function, and highlights the potent neuroprotective role of inhaled CBD. Using a commercially relevant nicotine delivery system, we demonstrate that while acute nicotine exposure transiently reduces seizure severity, chronic exposure significantly worsens seizure outcomes and induces coordinated molecular disruptions across multiple neurobiological systems. Critically, these detrimental effects are effectively reversed by inhaled CBD, underscoring its therapeutic potential in nicotine-related neurological dysfunction.

Behaviorally, we observed a biphasic response to nicotine exposure. Acute nicotine administration reduced seizure severity, likely via short-term modulation of cholinergic tone and suppression of excitatory signaling. In contrast, chronic exposure led to a marked escalation in seizure activity, mirroring maladaptive plasticity. This was paralleled by elevated expression of BDNF and c-FOS, markers of sustained neuronal activation and excitability. Notably, CBD inhalation reversed these molecular changes, restoring neuronal homeostasis under conditions of nicotine-induced stress.

A novel and central observation in this study is the disruption of glymphatic function following chronic nicotine exposure. We found a significant downregulation of Aquaporin-4 (AQP4), a key astrocytic water channel required for interstitial waste clearance. This represents, to our knowledge, the first evidence linking oral nicotine pouch use to impaired glymphatic signaling—a mechanism increasingly implicated in seizure susceptibility and neuroinflammatory burden. Remarkably, CBD inhalation restored AQP4 expression, suggesting that CBD may preserve or reestablish glymphatic integrity, offering a new therapeutic dimension in epilepsy management. Our data also reveal a tight association between seizure vulnerability and systemic inflammation. HMGB1 and IL-6, two hallmark biomarkers of peripheral and central immune activation, closely tracked the behavioral and molecular changes observed. While reduced under acute nicotine conditions, both markers were significantly elevated with chronic exposure. These increases likely reflect persistent inflammatory priming and neuroimmune crosstalk, contributing to heightened seizure risk. Importantly, CBD reversed these signatures, reaffirming its capacity to modulate not only neuronal, but also immune-driven aspects of seizure pathogenesis.

To integrate these findings, we employed a protein–protein interaction (PPI) network analysis using experimentally informed targets. The resulting nicotine-responsive network centered on IL-6, with interconnected nodes including FOS, BDNF, HMGB1, AQP4, and cholinergic regulators like Chrna7 and Lynx1. Gene Ontology (GO) enrichment revealed overrepresentation of processes related to IL-6 signaling, leukocyte differentiation, and inflammatory response. These systems-level insights highlight how chronic nicotine exposure destabilizes a multiscale molecular network spanning excitability, immune regulation, and fluid homeostasis. CBD’s ability to restore function across these domains suggests it acts as a global homeostatic regulator.

Altogether, our findings support a model in which chronic nicotine exposure promotes seizure susceptibility through dysregulation of interconnected excitatory, inflammatory, and glymphatic systems. CBD restores network stability by targeting critical nodes within this system, particularly those linked to IL-6-driven neuroimmune activation. These results offer a mechanistic foundation for the use of CBD as a therapeutic intervention in seizure disorders precipitated or exacerbated by nicotine exposure.

## Conclusion

This study provides the first integrated preclinical evidence that chronic exposure to oral nicotine pouches exacerbates seizure susceptibility, disrupts glymphatic function, and elevates key neuroinflammatory markers including HMGB1 and IL-6. In contrast, acute nicotine exposure exerts transient protective effects, emphasizing the complex, time-dependent nature of nicotine’s impact on brain health. Most importantly, our findings show that inhaled cannabidiol (CBD) robustly reverses these pathological effects—restoring neuronal excitability, immune homeostasis, and glymphatic integrity. Through combined behavioral, molecular, and systems biology approaches, we identify a nicotine-responsive network modulated by CBD, offering mechanistic insight into its neuroprotective actions. These findings strongly support the development of CBD inhalation as a non-invasive, targeted therapy for nicotine-related neurological complications, particularly in individuals at elevated risk for seizures, including chronic users of nicotine pouches.

## Author contributions

LPW, and BB: Study conception and design, analysis and interpretation of data, and drafting the article., BIB, ELS, SEN, AHA, and HMR: Acquisition of data and drafting the article., NY, JCY, MS, LPW and BB: Editing and Scientific Contribution.,

## Declaration of AI Assistance

AI-based software (e.g., ChatGPT) was used solely for language refinement and grammar editing; all scientific content and interpretation were entirely developed by the authors.

## Conflict of Interest

(1) Lei Phillip Wang, Babak Baban, and Jack Yu are members of Medicinal Cannabis of Georgia. (2) All other authors declare no conflict of interest. (3)Thriftmaster Holding Group (THG) is the provider of CBD inhalers and has a licensing contract with Augusta University. (4) THG had no role in study design, data collection and analysis, decision to publish, or preparation of the manuscript.

## Funding sources

This work was supported by institutional seed funding from the Dental College of Georgia at Augusta University.

## Acknowledgements

Authors are thankful to Thriftmaster Holding Group for providing the inhalant CBD for this study. Authors also thank Medicinal Cannabis of Georgia for providing help in the analysis, processing the data and optimizing the CBD dosage.

